# Increased cognitive complexity reveals abnormal brain network activity in individuals with corpus callosum dysgenesis

**DOI:** 10.1101/312629

**Authors:** Luke J. Hearne, Ryan J. Dean, Gail A. Robinson, Linda J. Richards, Jason B. Mattingley, Luca Cocchi

**Author notes:** <u>Corresponding author:</u> Luca Cocchi, QIMR Berghofer Medical Research, 300 Herston Road, Brisbane, QLD, 4006. Shared last author (Jason B. Mattingley, Luca Cocchi).

## Abstract

Cognitive reasoning is thought to require functional interactions between whole-brain networks. Such networks rely on both cerebral hemispheres, with the corpus callosum providing cross-hemispheric communication. Here we used high-field functional magnetic resonance imaging (7T fMRI), a well validated cognitive task, and brain network analyses to investigate the functional networks underlying cognitive reasoning in individuals with corpus callosum dysgenesis (CCD), an anatomical abnormality that affects the corpus callosum. Participants with CCD were asked to solve cognitive reasoning problems while their brain activity was measured using fMRI. The complexity of these problems was parametrically varied by changing the complexity of relations that needed to be established between shapes within each problem matrix. Behaviorally, participants showed a typical reduction in task performance as problem complexity increased. Task-evoked neural activity was observed in brain regions known to constitute two key cognitive control systems: the fronto-parietal and cingulo-opercular networks. Under low complexity demands, network topology and the patterns of local neural activity in the CCD group closely resembled those observed in neurotypical controls. By contrast, when asked to solve more complex problems, participants with CCD showed a reduction in neural activity and connectivity within the fronto-parietal network. These complexity-induced, as opposed to resting-state, differences in functional network activity help resolve the apparent paradox between preserved network architecture found at rest in CCD individuals, and the heterogeneous deficits they display in response to cognitive task demands [preprint: https://doi.org/10.1101/312629].

## Introduction

The corpus callosum is the major white matter commissure of the brain. It connects the left and right cerebral hemispheres, and consists of more than 190 million axonal projections (Tomasch, 1954). These connections are thought to support the coupling of bilateral functional networks (Honey et al., 2009; Putnam, Wig, Grafton, Kelley, & Gazzaniga, 2008; Roland et al., 2017; Shen et al., 2015; van der Knaap & van der Ham, 2011) that underpin both simple and complex behavior (Cocchi et al., 2014; Duncan, 2010; Schulte & Müller-Oehring, 2010). For example, adults who have had their corpus callosum surgically sectioned to control intractable seizures often experience a ‘disconnection syndrome’ in which they fail to complete tasks that require information-sharing between the two cerebral hemispheres (Gazzaniga, Bogen, & Sperry, 1962; Seymour, Reuter-lorenz, & Gazzaniga, 1994). By contrast, in individuals with *corpus callosum dysgenesis* (CCD), a neurodevelopmental condition in which the corpus callosum is only partially formed or where callosal axons fail to cross the cerebral midline (complete agenesis), deficits in behavioral measures of interhemispheric communication are generally more subtle (Lassonde, Sauerwein, Chicoine, & Geoffroy, 1991; Paul et al., 2007).

The neuropsychological profile of those with CCD are heterogeneous, ranging from subtle cognitive deficits to severe impairments across a wide variety of cognitive domains (Paul, Erickson, Hartman, & Brown, 2016; Paul, Schieffer, & Brown, 2004; Romaniello et al., 2017; Siffredi, Anderson, Leventer, & Spencer-Smith, 2013). Investigations using functional magnetic resonance imaging (fMRI) have reported intact resting-state functional brain networks in individuals with CCD (e.g., the default-mode network) (Owen et al., 2013; Tyszka, Kennedy, Adolphs, & Paul, 2011). Results from these studies suggest that the coupling of these networks is supported by alternative interhemispheric white matter pathways, such as those projecting through the anterior and posterior commissures (Lassonde et al., 1991; Tovar-Moll et al., 2014). However, such routes are indirect, polysynaptic and functionally costly (Marco et al., 2012; Owen et al., 2013).

The processing limits of functional networks shaped by CCD development are not well understood. Studies suggest that brain activity within sensory-motor networks during tactile stimulation and finger tapping tasks is preserved in CCD (Duquette, Rainville, Alary, Lassonde, & Lepore, 2008; Quigley et al., 2003). However, less is known about the function of brain networks associated with higher cognitive abilities (e.g., language; Hinkley et al., 2016). In neurotypical individuals, the execution of complex cognitive tasks is associated with increased activity within, and coupling between, fronto-parietal (FPN) and cingulo-opercular networks (CON) (Cocchi et al., 2014; Crittenden, Mitchell, & Duncan, 2016; Duncan, 2010). Whether individuals with CCD produce similar task-evoked patterns of brain network activity, and to what degree indirect pathways can support functional network coupling as task demands increase, is not currently known. It is likely that normal brain network topology and activity will be detectable in states of low or absent cognitive load (e.g., in the resting-state) (Owen et al., 2013; Tovar-Moll et al., 2014; Tyszka et al., 2011). By contrast, altered topology and deficits in network activity may emerge as cognitive demands increase, revealing functional limits in brain network plasticity.

To test this hypothesis, we used high field (7T) fMRI to characterize functional brain networks in seven individuals with CCD. We looked for changes in network configuration as individuals engaged in a well validated test of cognitive reasoning called the Latin Square Task (LST) (Birney, Halford, & Andrews, 2006). The LST is a non-verbal relational reasoning paradigm, similar to the popular game of Sudoku in its construction and rules, in which task complexity is parametrically varied with minimal demands on working memory (Birney et al., 2006; Halford, Wilson, & Phillips, 1998). We compared brain activation and network responses obtained in the seven CCD individuals with those from a normative sample of 30 neurologically typical participants. We identified specific functional connectivity deficits in CCD individuals, which only emerged under conditions of high cognitive demand.

## Materials and Methods

### Participants

Participants were recruited prospectively between October 2014 to April 2016. Individuals with CCD enrolled in the Australian database of corpus callosum disorders. CCD participants resided in, or travelled to, the city of Brisbane to undertake cognitive assessments and high-field magnetic resonance imaging (7T MRI) scans. The current study included seven participants with CCD aged 24–64 years (two females, one left handed participant, one ambidextrous) with levels of intelligence within the average range or above [Wechsler Abbreviated Scale of Intelligence, Full Scale IQ 88-124, mean = 102.29, SD = 13.35; see

**Table 1** for scores on additional measures of fluid (Advanced Raven’s Progressive Matrices) and crystalized intelligence (National Adult Reading Test)]. Participants CCD1, CCD2 and CCD4 had complete agenesis of the corpus callosum, whereas the remaining individuals had partial agenesis (see **Figure 1A** for a description of the corpus callosum abnormality). In line with previous work (Tovar-Moll et al., 2014), the CCD sample was heterogeneous, with different degrees of inter- and intra-hemispheric structural connectivity (**Figure 1B**, see methods below).

**Table 1.**
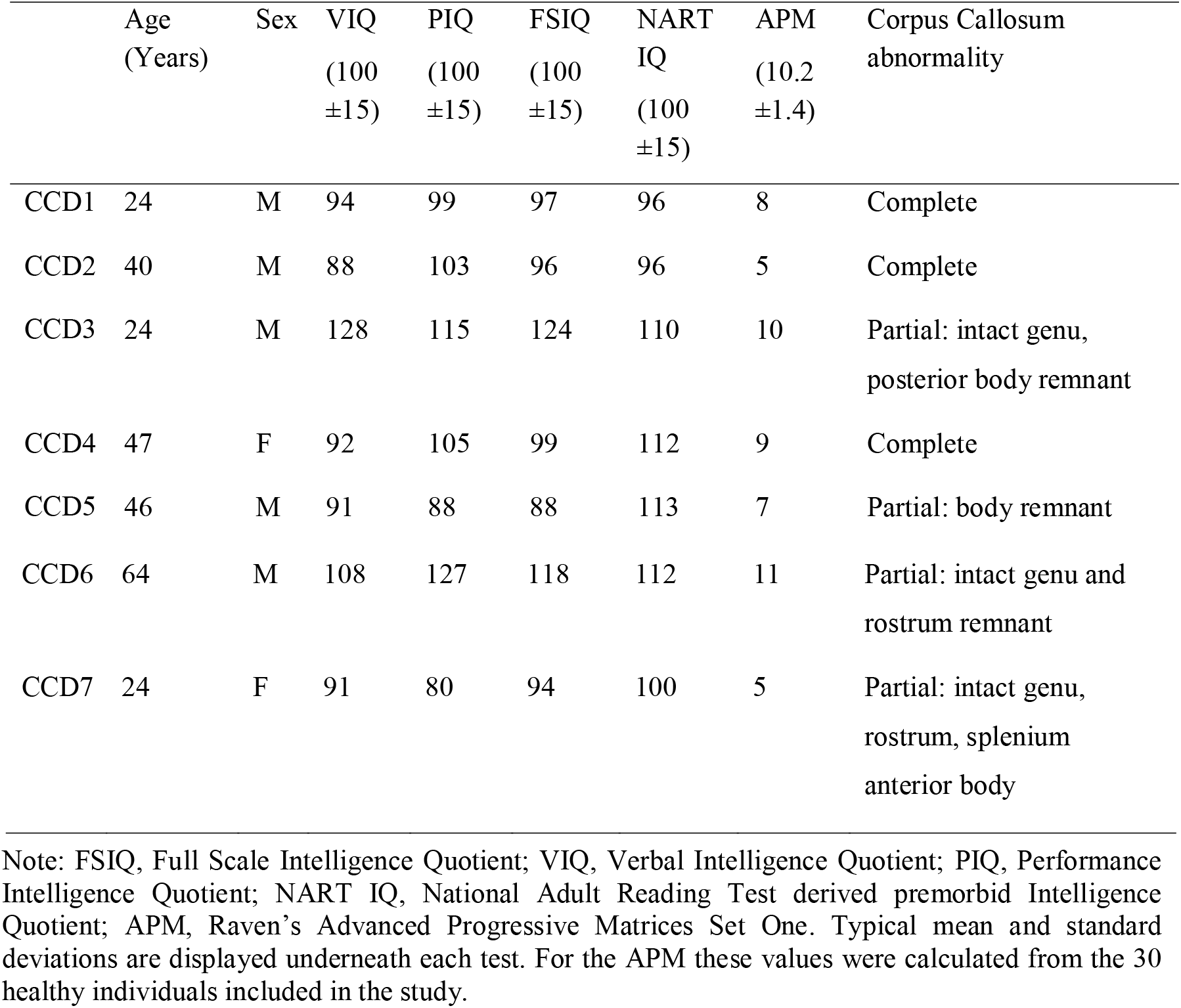
Demographics of the CCD participants and their scores on intellectual function measures.

**Figure 1.**
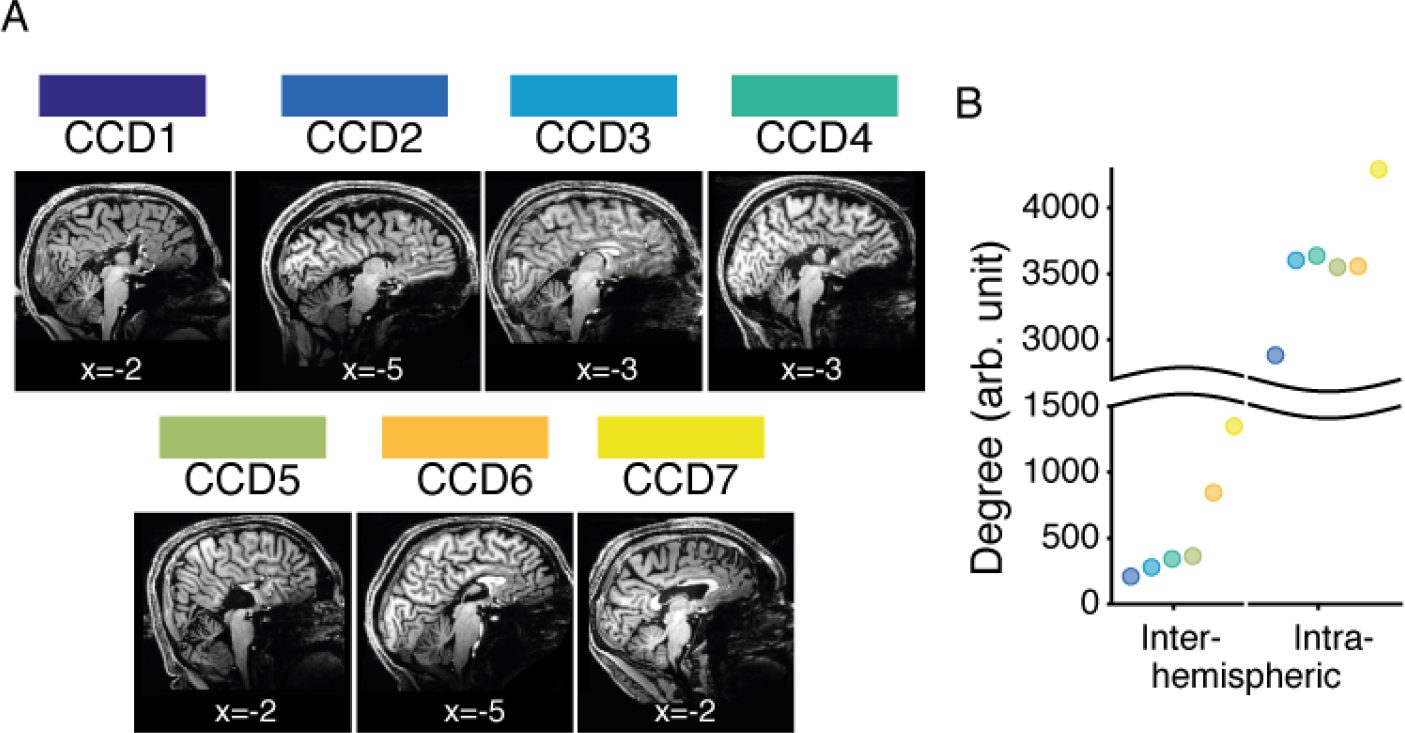
Structural anomalies in the seven corpus callosum dysgenesis (CCD) participants. **A.** Sagittal view (Montreal Neurological Institute, MNI, x coordinate indicated) of midline brain in individuals with CCD. Participants CCD1, CCD2 and CCD4 had complete agenesis of the corpus callosum. The other four participants had partial dysgenesis of the corpus callous with intact remnants: CCD3 had a preserved genu and posterior body; CCD5 had only a preserved body; CCD6 had an intact genu and rostrum remnant; and CCD7 had an intact genu, rostrum and splenium, as well as the anterior portion of the body. **B.** The number of structural connections across (inter) and within (intra) hemispheres was summed across ipsilateral and contralateral paired regions in each connectome (N = 6, CCD2 - CCD7). Dots are color-coded according to the convention depicted in panel **A**. As expected, participants with complete agenesis of the corpus callosum had fewer interhemispheric white matter connections (indicated by smaller numbers on the *y axis*) than those with a partially formed callosum. This trend was similar for intra-hemispheric connections. DWI data for CCD1 were not available due to a technical problem with data acquisition.

To benchmark participants’ behavioral performance, their patterns of task-induced brain activity and their functional connectivity, we included a sample of 30 healthy controls (mean age = 23.60, SD = 3.95 years; 17 females, all right handed). The controls were a subsample of a previously published study (Hearne, Cocchi, Zalesky, & Mattingley, 2017) who had completed the same task and imaging protocols. Individuals with the most comparable gender, age and behavioral performance profiles to the CCD individuals were included in the current cohort. These participants were not administered the neuropsychological battery (Table 1) but were compared in terms of their behavioural performance, fluid intelligence (Raven’s Advanced Progressive Matrices), gender, handedness and age.

All methods and procedures were approved by The University of Queensland Human Research Ethics Committee and written informed consent was obtained from all participants.

### Experimental design and statistical analysis

Participants completed three × 12-minute runs of the Latin Square Task (LST) (Birney et al., 2006), high-angular resolution diffusion MRI, and an MP2RAGE anatomical scan.

#### Task

The LST was identical to the task used in our previous work (Hearne et al., 2017). Each trial involved the presentation of a 4 × 4 matrix populated with a small number of geometric shapes (square, circle, triangle or cross), blank spaces and a single target question mark (‘?’, see **Figure 2A**). Participants were asked to solve for the cell containing the question mark, adhering to the rule: “*each shape can only occur once in every row and once in every column”.* Binary problems require integration of information across a single row *or* column. Ternary problems involve integration across a single row *and* column. Quaternary problems require integration of information across several rows and columns. Null trials involved presentation of an LST grid, but instead of a target probe (‘?’) an asterisk was presented (‘*’) indicating to the participant that no reasoning was required. For the brain imaging analyses, the Null trials were utilized as ‘baseline’ comparison periods (see analysis methods below). Puzzles were presented for five seconds, followed by a two second motor response period and confidence rating (**Figure 2B**). In total, 144 LST items were presented in the MRI session across 16 blocks, with 36 items in each condition. Trials were pseudo-randomized such that no two items of the same level of complexity occurred sequentially, and each block had two problems from each level of complexity. Participants were not cued to the complexity of any given trial. Both groups were provided with the same task instructions, and prior to the MRI session all participants completed 20 practice trials of the LST (12 with corrective feedback).

**Figure 2.**
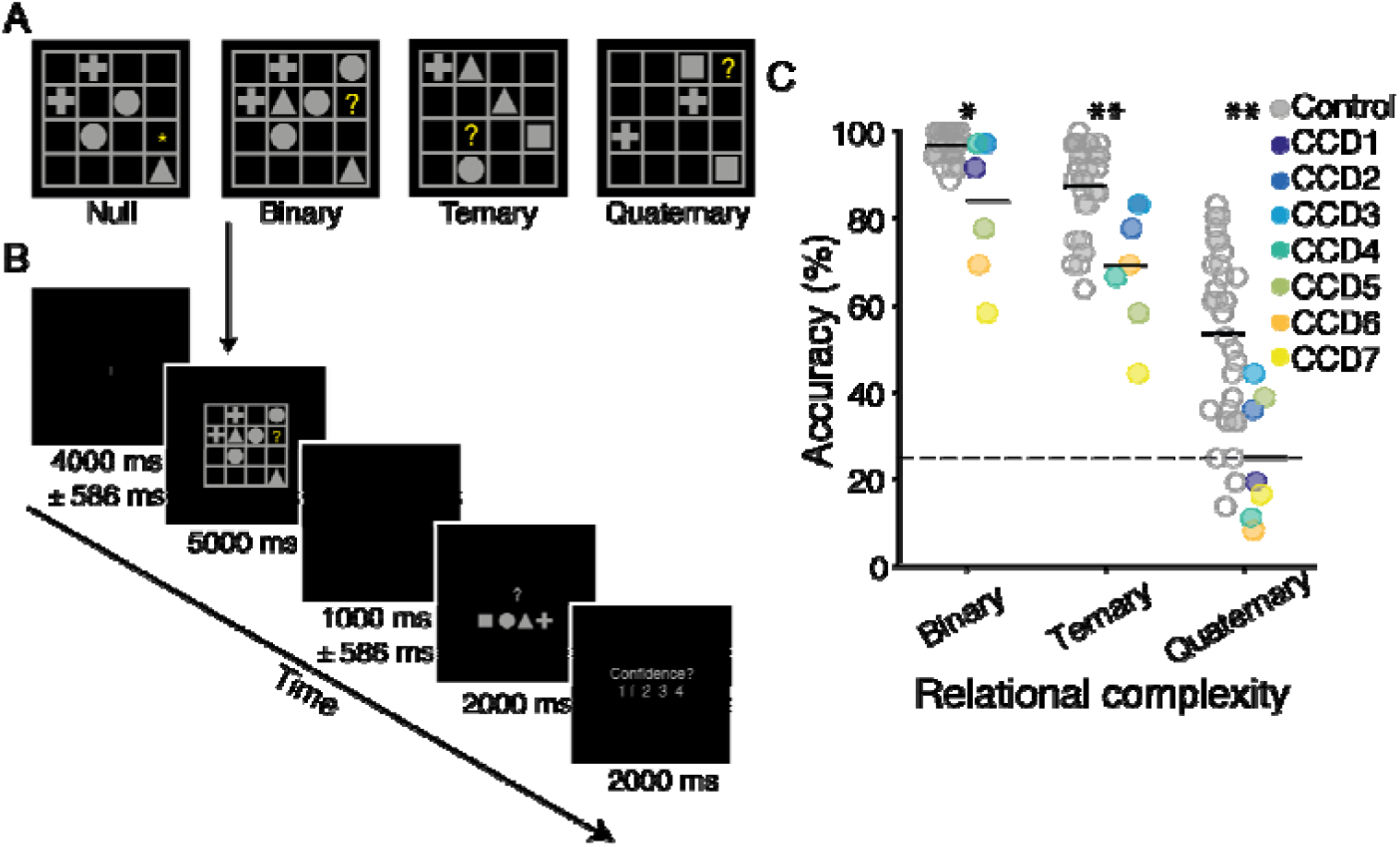
Latin Square Task design and behavioral performance of individuals with corpus callosum dysgenesis (N=7) and controls (N=30). **A.** Examples of each reasoning complexity condition. The correct answers are square, cross and cross, respectively, for the binary, ternary and quaternary problems illustrated. On null trials no reasoning was required. **B.** Example trial sequence. Each trial contained a jittered fixation period, followed by an LST item, a second, jittered fixation period, a response screen, and a confidence rating scale. In null trials the motor response screen had one geometric shape replaced with an asterisk, which indicated the button to press for that trial. **C.** LST performance accuracy as a function of reasoning complexity for the two groups (controls on the left and CCD group on the right). Values for each individual are plotted for the controls (gray) and CCD individuals (colors). Black horizontal line-segments represent the mean value for each group and condition. ^*^ indicates *p* < .05, ^**^ indicates *p* < .01; Wilcoxon rank sum test.

#### Neuroimaging data acquisition

Imaging data for all participants were acquired using a 7 Tesla Siemens Magnetom whole body MRI scanner fitted with a 32-channel head coil. Structural (T1), functional (fMRI), and diffusion-weighted (DWI) MRI was conducted for each participant.

Details regarding the fMRI protocol have been reported elsewhere (Hearne et al., 2017). Briefly, whole brain T2*-weighted gradient echo images were acquired using an optimized multi-band sequence (acceleration factor of five). In each task run, 1250 volumes were collected with the following parameters: voxel size = 2 × 2 × 2 mm, TR = 586 ms, TE = 23 ms, flip angle = 40°, FOV 121 × 208 × 208 mm, 55 slices.

The structural MRI protocol used an MP2RAGE sequence (voxel size = 0.75 × 0.75 × 0.75 mm, MTX 256 × 300 × 320, TR = 4300 ms, TE = 3.44 ms, TI = 840 / 2370 ms, flip angle 5°, FOV 192 × 225 × 240 mm). The DWI protocol used a stimulated-echo diffusion-weighted echo planar imaging (EPI) sequence (voxel size = 1.55 × 1.55 × 1.55, MTX 142 × 142 × 70, FOV 220 × 220 × 126 mm (0.3 mm axial slice gap), Δ / δ = 110 / 30 ms, TR = 12000 ms, TE = 54.4 ms). Diffusion space was evenly sampled over two half-shells (32 sample points with *b* = 700 s mm^-2^ and 64 sample points with *b* = 2000 s mm^-2^, respectively). Diffusion gradient directions were generated using an electrostatic energy minimization method. Four *B*_0_ (i.e., *b* ≈ 0 s mm^-2^) images were also acquired (one of which was reverse phase-encoded to correct EPI associated image distortions). DWI data for CCD1 were not available due to a technical problem with the acquisition sequence.

#### Pre-processing

Functional imaging data were pre-processed using the MATLAB (Mathworks, USA) toolbox Data Processing Assistant for Resting-State fMRI (DPARSF V 3.0, Chao-Gan and Yu-Feng, 2010). DICOM images were first converted to Nifti format and realigned. T1 images were re-oriented, skull-stripped and co-registered to the Nifti functional images. Segmentation (gray matter, white matter and cerebrospinal fluid) and the DARTEL algorithm (Ashburner, 2007) were used to improve the estimation of non-neural signal in subject space. From each gray matter voxel, the following signals were regressed: undesired linear trends, signals from the six head motion parameters (three translation, three rotation), white matter and cerebrospinal fluid (estimated from individual-level seed regions within the CSF and white matter). Single-subject functional images were then normalized and smoothed using DARTEL (4 mm^3^). Data processing steps also involved filtering (0.01–0.15 Hz) at a low frequency component of the blood oxygenation level dependent (BOLD) signal known to be sensitive to both resting state and task-based neural activity (Bassett, Yang, Wymbs, & Grafton, 2015; Braun et al., 2012; Sun, Miller, & D’Esposito, 2004).

The DWI data were processed using MRtrix3 (www.mrtrix.org). The diffusion-weighted images were corrected for EPI distortion, motion, and bias field inhomogeneity before the structural images were registered to the diffusion-weighted images. The co-registered structural images were then decomposed into five tissue type (5TT) images: Cortical gray matter, sub-cortical gray matter, white matter, CSF, and pathological tissue. The 5TT images were used in conjunction with the DWI data to create separate fiber orientation distribution function (fODF) maps for WM, grey matter (GM) and cerebrospinal fluid (CSF) via the multi-shell, multi-tissue, constrained spherical devolution (MSMT-CSD) algorithm, with a maximum spherical harmonic order of eight (Dhollander, Raffelt, & Connelly, 2016; Jeurissen, Tournier, Dhollander, Connelly, & Sijbers, 2014).

#### Task-evoked activity

The mean performance of CCD individuals in the Quaternary condition (**Figure 2C**) was at chance level. We therefore excluded data from this condition for the imaging analyses. Regional activity changes associated with increments in relational complexity were isolated using the general linear model (GLM) framework implemented in SPM12 (Friston et al., 1994). Fixation, reasoning (Null, Binary, Ternary), motor response, and confidence rating epochs were each modelled as boxcar functions convolved with a canonical hemodynamic response function. Temporal autocorrelations were corrected using the FAST algorithm due to the short TR (Bollmann, Puckett, Cunnington, & Barth, 2018).

Single subject (first-level) *t*-contrasts were used to isolate the average positive effect of increased relational complexity [Null (-2) *versus* Binary (1) and Ternary (1)]. In the control group, first level contrasts were used to undertake a one-sample *t*-test to isolate the average positive task effect at group-level. An initial uncorrected threshold of *p* < 0.0001 was applied to define clusters of brain regions. Cluster level correction was subsequently ascribed to declare significance [p < 0.001, family-wise error corrected (FWE)]. Considering the small sample size of the CCD cohort, we generated binarized activity maps containing the highest *t*-statistics for each individual (top 5% of values at voxel level). These maps were then overlaid to create a group-level ‘consistency map’. The rationale behind these analyses was to isolate brain clusters involved in LST performance in both groups. Results from the two groups were qualitatively compared (**Results** section). The SPM toolbox MarsBar (Brett, Anton, Valabregue, & Poline, 2002) was subsequently used to extract single-subject beta-values from twelve regions of interest (ROI, 5mm spheres; **Table 2**). The ROI were defined based on fMRI activations (see **Results**) and were given brain network affiliations in line with previous literature (Cocchi et al., 2014; Cole & Schneider, 2007; Dosenbach et al., 2006; Duncan, 2010). Six ROI mapped onto the fronto-parietal network (FPN) and six onto the cingulo-opercular network (CON). The placement of each individual ROI was manually checked to confirm that differences in neuroanatomy within the CCD cohort did not lead to sampling error. To test for changes in neural activity between groups and reasoning complexity, mean beta values for each network were entered into a two (group) by two [complexity (Null versus Binary, Null versus Ternary)] mixed ANOVAs.

**Table 2.**
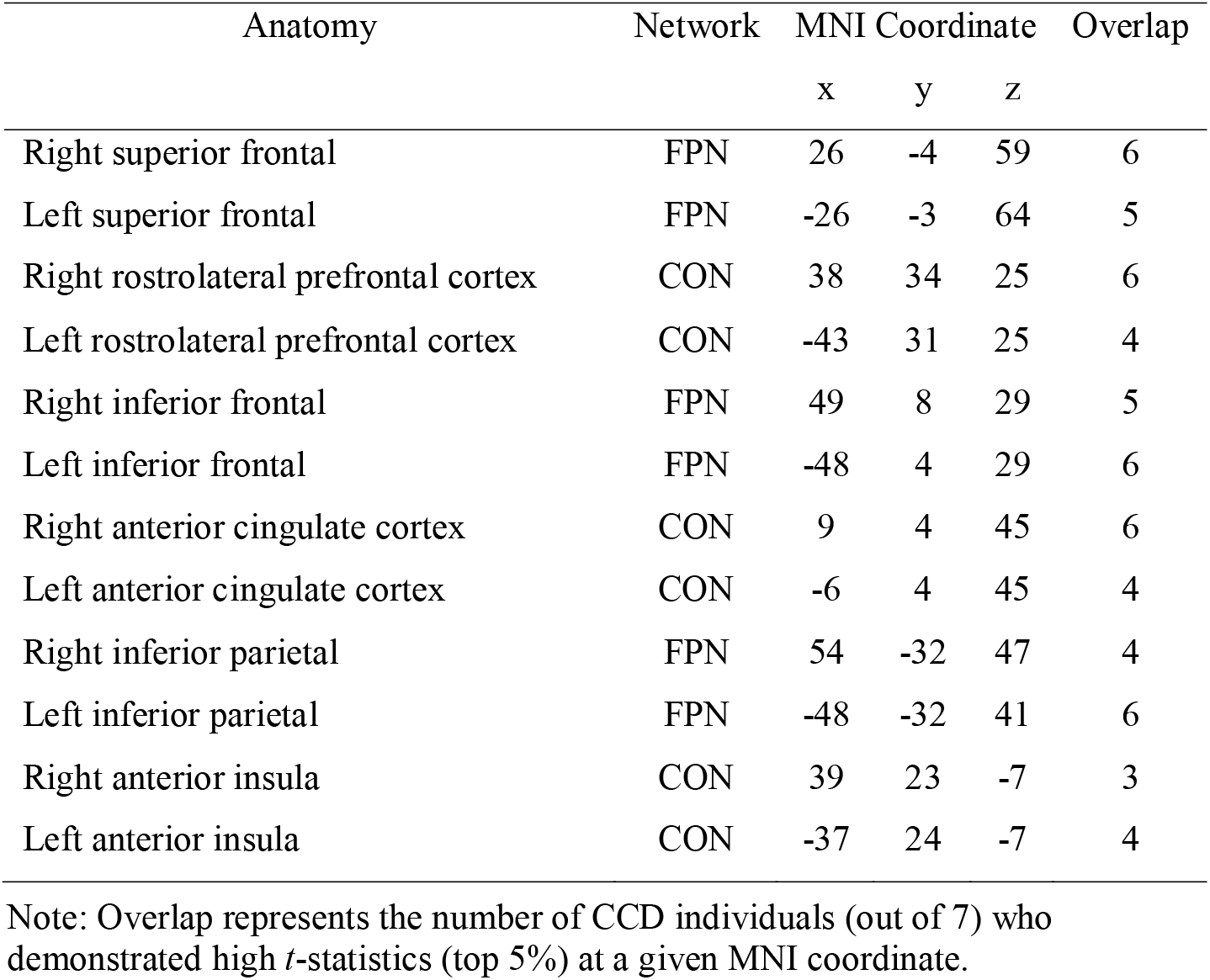
Regions of interest derived from the GLM analysis.

#### Task-evoked functional connectivity

Task-evoked functional connectivity was estimated using the same method adopted in Hearne et al., (Hearne et al., 2017). In short, regionally averaged time-series were extracted for the twelve ROIs, and a task nuisance covariate was regressed from each signal (Cole, Bassett, Power, Braver, & Petersen, 2014). After accounting for the hemodynamic lag, the residual values were concatenated to form a condition-specific (Null, Binary, Ternary) time-series of interest. A Pearson correlation was performed on the residual time-series. Connectivity matrices were fisher-Z transformed to improve the use of parametric statistics. To test for differences in functional connectivity across complexity conditions and groups we conducted two (group) by three (complexity) mixed ANOVAs on mean connectivity values in each network on both inter- and intra-hemispheric connections.

#### Structural connectivity

Structural connectivity analyses were undertaken using MRtrix3 (Tournier, Calamante, & Connelly, 2012). First, CSD-based probabilistic tractography (using the iFOD2 algorithm) (Tournier, Calamante, & Connelly, 2010) was used to generate whole-brain tractograms for each participant. The tractograms consisted of 20 million streamlines each that were seeded randomly across the whole-brain white matter. The streamlines were anatomically constrained to the WM to ensure streamlines did not path through GM, CSF, or potential lesions (Smith, Tournier, Calamante, & Connelly, 2012). The tracking parameters were: fODF cut-off = 0.1, step size = 0.5 mm, curvature threshold = 45°/step, minimum / maximum streamline length = 10 / 200 mm. These tractograms were then parsed by a CSD-informed tractogram filter (Smith, Tournier, Calamante, & Connelly, 2013) to create a new tractogram consisting of the 2 million streamlines that represented the most likely propagations of fiber tracts. Binary connectomes (matrix representations of the brain network topology) were generated from the filtered tractograms using the Harvard-Oxford cortical and subcortical structural parcellation atlas (Desikan et al., 2006; Frazier et al., 2005; Goldstein et al., 2007; Makris et al., 2006). The atlas was warped into the native space for each set of diffusion-weighted images by co-registering the associated MNI 152 template to these images. To estimate the extent of inter- and intra-hemispheric structural connectivity, the number of connections (i) within and (ii) across hemispheres was summed across ipsilateral and contralateral paired regions in each connectome (**Figure 1B**). Further analysis taking connection weight into account showed similar results.

## Results

### Behavior

For the LST, a typical complexity effect was observed in individuals with CCD, as previously found in independent samples of neurotypical adults (Birney & Bowman, 2009; Cocchi et al., 2014) (**Figure 2C**). Reasoning problems of greater complexity resulted in significantly more reasoning errors (*F*_2,12_ = 60.45, *p* < .001, *n*^2^_*p*_ = .91) and longer reaction times (RT; *F*_2,12_ = 6.31, *p* = .013, *n*^2^_*p*_ = .51). Follow-up tests confirmed that participants were generally more accurate, and faster, in the Binary than in the Ternary condition (accuracy: *t*_6_ = 4.15, *p* = .006, *d* = 1.57, RT: *t*_6_ = 3.89, *p* = .008, *d* = 1.47), and in the Ternary than in the Quaternary condition (accuracy: *t*_6_ = 6.89, *p* < .001, *d* = 2.60; RT: n.s., *t*_6_ = 0.67, *p* = .53, *d* = 0.25) (**Figure 2C**).

When compared with controls, a mixed ANOVA (complexity × group) revealed a significant main effect of group for accuracy (*F*_2,70_ = 27.26, *p* < .001, *n*^2^_*p*_ = .44) and RT (*F*_2,70_ = 9.27, *p* = .004, *n*^2^_*p*_ = .21), such that controls were faster and more accurate than CCD individuals. The CCD cohort showed reduced accuracy and slower reaction times compared to controls at all complexity levels (*p* < .05). No interaction was observed for either accuracy or RT (**Figure 2C**). CCD individuals also showed reduced performance on the APM (M = 7.86, SD = 2.34) compared with controls (M = 10.17, SD = 1.37, *z* = 2.47, *p* = .01), which is consistent with previous work suggesting a relationship between the LST and fluid intelligence (Birney & Bowman, 2009; Hearne et al., 2017). Overall, the behavioral results suggest that CCD individuals were slower and more error-prone than controls but showed an analogous complexity-induced decline in accuracy and increase in response speed. Four CCD individuals performed at chance level in the Quaternary trials (**Figure 2C**), and so this condition was eliminated from the imaging analyses.

### Brain activity

We first tested whether individuals with CCD showed similar brain activity patterns as the control group in response to reasoning complexity. Previous work on cognitive reasoning has isolated co-activations within fronto-parietal (FPN) and cingulo-opercular networks (CON) that respond to task complexity, such that increased complexity is coupled with an increase in the BOLD response (Cocchi et al., 2014; Cole & Schneider, 2007; Dosenbach et al., 2006; Duncan, 2010). We calculated the average positive effect of increased relational complexity by comparing Null versus Binary and Ternary conditions in a general linear model (see **Methods** for details). Due to the small sample size of the CCD cohort, we generated binarized activity maps containing the highest *t*-statistics for each individual (voxel level, top 5% of values; range 0.7 - 27.5). These maps were then overlaid to create a ‘consistency map’ (shown in **Figure 3A**). Qualitatively, the resulting maps suggest that both groups share a similar pattern of neural co-activations in response to reasoning task demands, involving key cortical clusters in regions of the FPN and CON.

**Figure 3.**
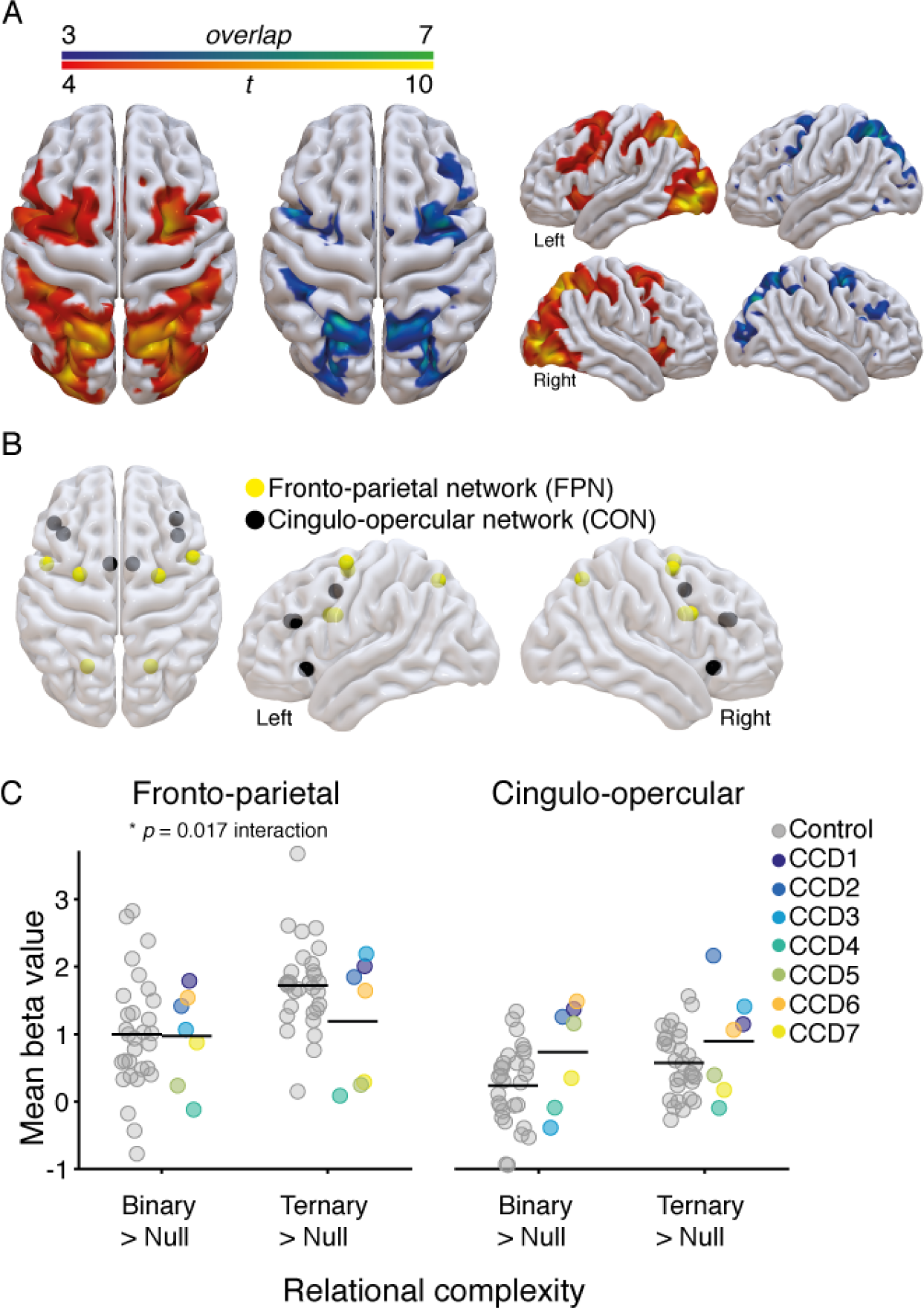
Brain activity associated with increases in reasoning complexity. **A.** Brain activity evoked in the CCD group (*N* = 7), comparing null versus binary and ternary conditions, contrasted with brain activity evoked in the control group (*N* = 30). The blue-green heat map (left) represents the overlap of brain activity across CCD individuals (voxel level top 5% of *t*-values), such that green represents complete overlap (100%, *N* = 7) and blue represents less overlap (42%, *N* = 3). The red-yellow heat map (right) shows the results of a random effects analysis in the control group (conservative threshold of *p* < 0.0001 uncorrected, p < 0.001 FWE corrected at cluster level). This stringent statistical threshold was adopted to highlight the key neural clusters involved in the task. Brain renderings were created using SurfIce software (www.nitrc.org/projects/surfice). **B.** Regions of interest (ROIs) plotted as spheres in a glass brain (details in **Table 2**). These brain regions overlap with the complexity-induced patterns of activity shown by the CCD and control participants, and comprise two well-known brain networks involved in cognitive control: the fronto-parietal (FPN) and cingulo-opercular (CON) networks (Cocchi et al., 2014; Dosenbach et al., 2006; Duncan, 2010; Seeley et al., 2007; Yeo et al., 2011). **C.** Change in the estimated beta values averaged across ROIs within the FPN (left) and CON (right). Values for each individual are plotted for the control (gray) and CCD cohort (color, legend in center). Black horizontal line-segments represent the mean value for each group and condition.

To investigate the magnitude of brain activation across conditions, we compared change in neural activity (beta values) within twelve regions of interest that were selected based upon the identified patterns of activity (**Figure 3B & Table 2).** A mixed ANOVA (complexity × group) on mean change in beta values across each brain network revealed a significant main effect of complexity, such that the Ternary condition yielded a larger increase in neural activity than the Binary condition when compared with the Null condition (FPN: *F*_1,35_ = 21.45, *p* < .001, *n*^2^_*p*_ = .38; CON: *F*_1,35_ = 6.2, *p* = .02, *n*^2^_*p*_ = .15; **Figure 3C**). For the FPN, a significant complexity by group interaction was observed (*F*_1,35_ = 6.30, *p* = .017, *n*^2^_*p*_ = .15). This interaction suggests that, contrary to controls, individuals with CCD showed a diminished increase in neural activity from Binary to Ternary conditions within the FPN (**Figure 3C**).

### Brain connectivity

Having characterized changes in complexity-induced brain activity, we next tested whether individuals with CCD demonstrated similar patterns of inter- and intra-hemispheric task-based functional connectivity to the control cohort in response to increasing relational complexity. Previous work has shown that increased relational complexity induces robust increases in functional connectivity between the FPN and CON (Cocchi et al., 2014). Task-based functional connectivity was calculated between regions that showed the most consistent BOLD activation across all participants (**Table 2**).

The CCD cohort had similar levels of mean functional connectivity to controls in the Null condition, supporting previous work showing intact bilateral networks in CCD individuals under low cognitive load (Owen et al., 2013; Tyszka et al., 2011). By contrast, a mixed ANOVA with factors of complexity and group revealed an interaction in the FPN (inter-hemispheric: *F*_2,70_ = 3.69, *p* = .030, *n*^2^_*p*_ = .10, intra-hemispheric: *F*_2,70_ = 3.37, *p* = .040, *n*^2^_*p*_ = .09, **Figure 4A**). In this network, CCD individuals showed a reduction in functional connectivity as a function of complexity compared with controls (**Figure 4A**).

**Figure 4.**
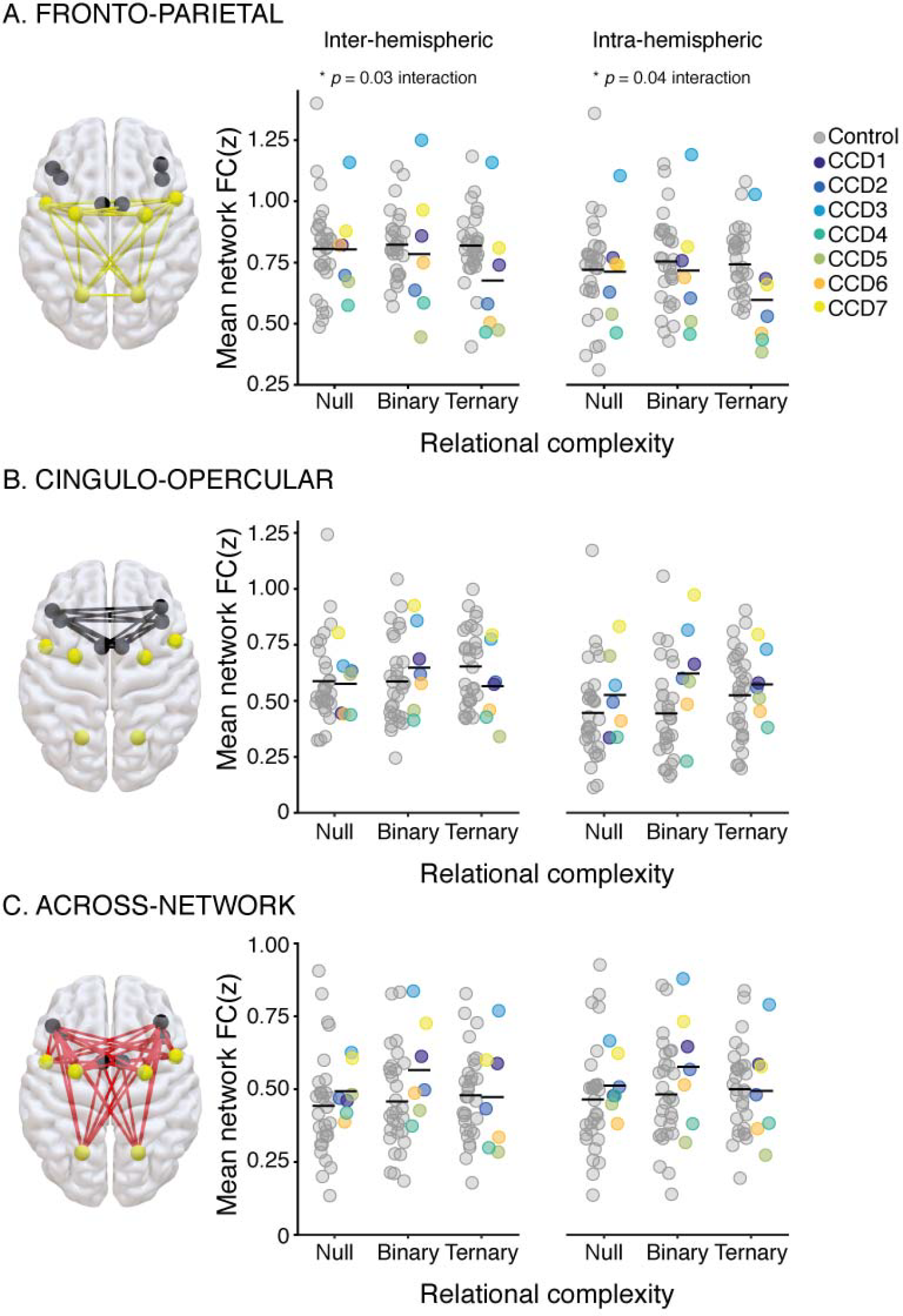
Within- and across-network functional connectivity associated with increasing reasoning complexity. Mean inter-hemispheric (left) and intra-hemispheric (right) *z*-transformed functional connectivity values (*y-axis*) plotted as a function of increasing reasoning complexity demands (*x-axis*: Null, Binary, Ternary). **A.** Fronto-Parietal (FPN) connections. **B.** Cingulo-Opercular (CON) connections. **C.** Across-network connections. Brain renderings of the included connections are on the right (yellow for FPN, black for CON, and red for across-network connections). Values for each individual are plotted for the controls (gray circles) and CCD individuals (colored dots, legend top middle). Black horizontal line-segments represent the mean value for each group and condition.

### Control analysis

A Wilcoxon rank sum test revealed differences in age between individuals with CCD and the control cohort (M_CCD_ = 38.86, SD_CCD_ = 15.06, M_HC_ = 22.29, SD_HC_ = 5.25, *z* = -3.36, *p* < .001). As a control analysis we plotted changes in task accuracy (Ternary – Binary) beta values (Ternary – Null) and functional connectivity (Ternary – Null) as a function of individuals’ age. Numerically, values from the oldest CCD participants were within the range of younger healthy controls. For example, in **Figure 5C** the oldest CCD participant shows the lowest connectivity value (z-value), while the next lowest value is from an 18-year-old healthy control. These analyses suggest that between group differences in age are unlikely to explain our findings.

**Figure 5.**
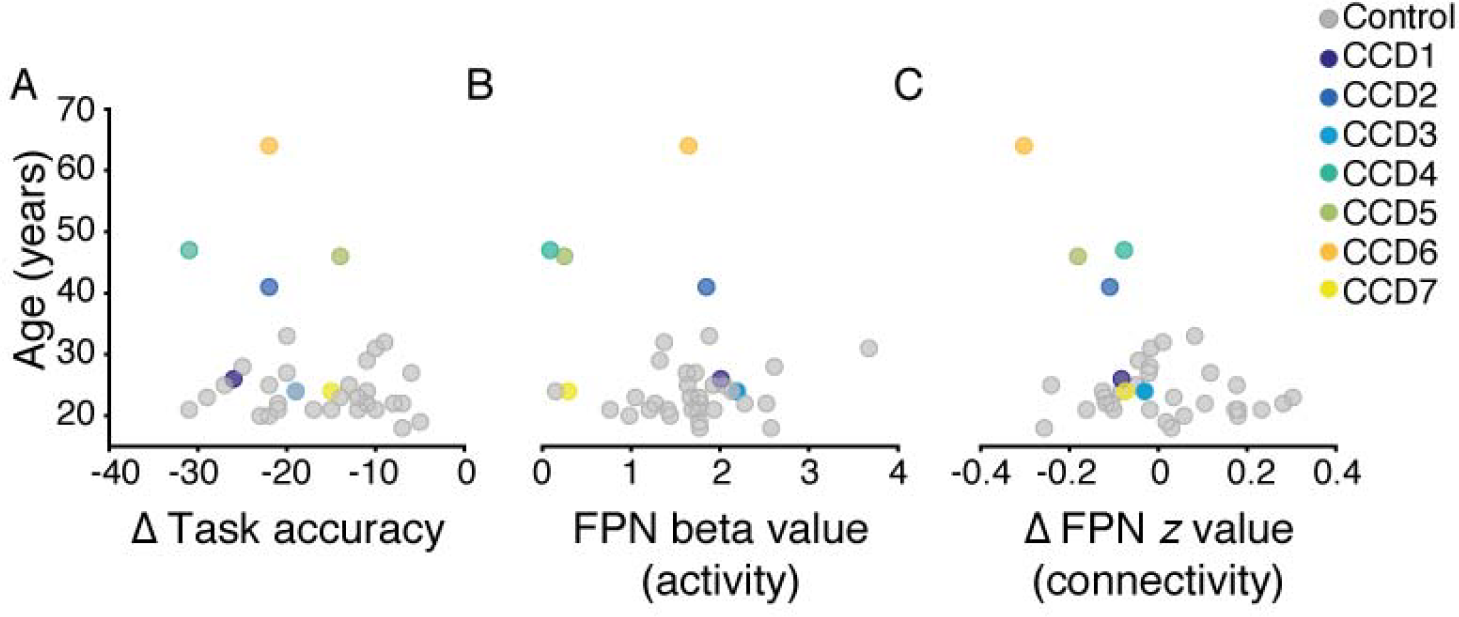
Control analyses to investigate the effect of age. Scatter plots of changes in task accuracy (**A**), FPN activity (**B**), and FPN connectivity (**C**) plotted as a function of participant age (*y*-axis).

## Discussion

Despite the absence or malformation of the corpus callosum, preserved bilateral functional brain networks have been observed in individuals with CCD (Owen et al., 2013; Tovar-Moll et al., 2014; Tyszka et al., 2011). These whole-brain functional networks are thought to be supported by atypical interhemispheric white matter pathways that form during development (Lassonde et al., 1991; Tovar-Moll et al., 2014). Less is known about the role and functional properties of brain networks supported by this abnormal connectome. Here we had individuals with CCD undergo fMRI while solving non-verbal reasoning problems known to elicit robust neural responses within and between the FPN and CON (Cocchi et al., 2014; Crittenden et al., 2016; Hearne, Cocchi, Zalesky, & Mattingley, 2015; Hearne et al., 2017). Behaviorally, individuals with CCD showed a typical complexity effect, in that their performance declined as a function of increased reasoning complexity (Birney & Bowman, 2009; Hearne et al., 2017). Neural activity and connectivity patterns within and between the FPN and CON networks in CCD individuals were largely preserved in low-complexity trials. By contrast, abnormal patterns of activity and connectivity emerged in this group under higher complexity conditions, specifically within the FPN. These findings reveal previously unknown state-dependent deficits of functional integration in individuals with CCD, which evidently remain hidden at rest or when cognitive demands are low.

Our results support the notion that resting-state functional networks underpinning higher cognitive processes can be supported by abnormal configurations of the connectome (Mišić et al., 2016; Owen et al., 2013; Tyszka et al., 2011). Under low complexity conditions (i.e., the Null or Binary problems of the LST), we observed relatively normal neural activity and patterns of functional connectivity within and between segregated brain regions of the FPN and CON in CCD individuals (**Figure 3** & **4**). Strikingly, even participants with complete agenesis of the corpus callosum showed bilateral patterns of activity and connectivity in these networks. Subcortical structures, as well as the anterior and posterior commissures (Tovar-Moll et al., 2014), can provide common input to both cerebral hemispheres, leading to observable bilateral functional neural networks. The thalamus, for example, is endowed with the type of whole-brain structural and functional connectivity necessary for such a role (Behrens et al., 2003; Bell & Shine, 2016; Hwang, Bertolero, Liu, & D’Esposito, 2017; Schmitt et al., 2017), as well as the potential to support direct inter-hemispheric communication through the interthalamic adhesion (Damle et al., 2017). Our findings in CCD individuals provide motivation for further work to assess the putative contribution of subcortical routes in supporting bilateral brain activity in pathologies that impact the corpus callosum.

Under conditions of high reasoning demands, however, we found that individuals with CCD showed decreased local neural activity and functional connectivity within the FPN. In terms of brain activity, individuals with CCD showed a smaller increase in neural activity compared to controls when attempting to solve more complex reasoning problems. Moreover, CCD individuals showed a decline in functional connectivity within the FPN as reasoning complexity increased, whereas controls instead showed a shift toward increased functional connectivity with increasing cognitive load across both the FPN and CON (**Figure 4**) (Cocchi et al., 2014). Critically, we found little distinction between intra- and inter-hemispheric connectivity in regards to this deficit, suggesting a global impact of CCD on task-induced FPN dynamics. The FPN is thought to be responsible for the moment-to-moment implementation of cognitive task goals (Cocchi, Zalesky, Fornito, & Mattingley, 2013; Dosenbach et al., 2007). Damage to frontal and parietal cortices has also been associated with poorer performance in tests of cognitive reasoning (Urbanski et al., 2016; Woolgar et al., 2010). In line with this, our results suggest that in CCD individuals FPN activity does not modulate in the manner required to solve complex cognitive problems.

CCD is a developmental malformation of the central nervous system, affecting approximately 1:4000 people (Glass, Shaw, Ma, & Sherr, 2008). The current study included seven individuals with CCD recruited from Australia over a period of 18 months. While this sample size is comparable to, or larger than, that of several previous investigations (Owen et al., 2013; Quigley et al., 2003; Tovar-Moll et al., 2007, 2014; Tyszka et al., 2011; Wahl et al., 2009), it was nevertheless too small for us to use many of the conventional analytic approaches for inferring differences in neural activity and connectivity between groups (Eklund, Nichols, & Knutsson, 2016; Friston et al., 1994; Zalesky, Fornito, & Bullmore, 2010). We therefore opted for an analysis strategy that balanced statistical rigor with the reality of describing whole-brain neural networks in a rare and heterogeneous condition. Group by complexity interactions for connectivity values did not quite survive a strict correction for multiple comparisons. However, altered complexity-induced changes in connectivity were consistent with the observed abnormalities in FPN activity. Our control analyses argue against a major impact of age on the results, but we cannot rule out a possible influence of this variable, or other potential confounding variables, such as differences in handedness, on the observed interaction. Likewise, CCD has been associated with additional heterogeneous cerebral malformations including, for example, abnormal sulcation, noncallosal midline abnormalities, and differences in axonal demyelination, that likely influence behavioural and functional imaging outcomes (Barkovich & Norman, 1988; Hetts, Sherr, Chao, Gobuty, & Barkovich, 2006; Parrish, Roessmann, & Levinsohn, 1979). Finally, due to the small sample size the possibility of more subtle and undetected deficits in complexity-induced network activity in CCD cannot be excluded. It is important to consider the current findings as initial evidence to guide further, more definitive research with larger sample sizes.

In conclusion, the current findings are in line with the notion that resting-state functional brain networks supporting cognitive control are at least partially preserved in individuals with CCD (Tyszka et al., 2011). This observation also provides an empirical demonstration of the brain’s ability to accommodate canonical functional networks within different anatomical scaffolds (Mišić et al., 2016). Despite preserved network activity and functional connectivity under low-complexity reasoning demands, the FPN in individuals with CCD did not seem to effectively accommodate increasingly complex task demands. This suggests that, even with alternative pathways to support brain activity, deficits in corpus callosum integrity impose a hard limit on the capacity of the FPN to dynamically adapt its activity to high cognitive demands. When considered in conjunction with individual differences in structural connectivity, task-driven deficits in FPN integration provide a plausible neural mechanism to explain the broad and heterogeneous cognitive deficits commonly observed in individuals with CCD.

## Acknowledgements

The authors thank Tim Edwards, Annalisa Paolina, and Sinead Eyre for assistance with participant recruitment and MRI data collection, Jacquelyn Knight and Megan Barker for compiling the neuropsychological data, and Dr Kieran O’Brien, Dr Markus Barth, and Dr Steffen Bollmann for MRI sequence optimizations. The authors acknowledge the facilities, and the scientific and technical assistance of the National Imaging Facility at the Centre for Advanced Imaging, University of Queensland. The authors are indebted to the participants for generously donating their time to be part of this study and thank the Australian Disorders of the Corpus Callosum (AusDoCC) support group for their assistance in recruiting CCD participants for the database and this study. This research was supported by the Australian Research Council (ARC) ARC-SRI Science of Learning Research Centre (SR120300015), and the ARC Centre of Excellence for Integrative Brain Function (ARC Centre Grant CE140100007). J.B.M was supported by an ARC Australian Laureate Fellowship (FL110100103). L.C. was supported by an NHMRC grant (APP1099082). G.A.R. was supported by an ARC DECRA Fellowship (DE120101119). L.J.H was supported by an Australian Postgraduate Award. R.J.D was supported by funds from Brain Injured Children’s Aftercare Recovery Endeavors (BICARE) and L.J.R. was supported by an NHMRC Principal Research Fellowship (APP1120615).

## References

Ashburner, J. (2007). A fast diffeomorphic image registration algorithm. NeuroImage, 38(1), 95–113. http://doi.org/http://dx.doi.org/10.1016/j.neuroimage.2007.07.007

Barkovich, A. J., & Norman, D. (1988). Anomalies of the corpus callosum: correlation with further anomalies of the brain. American Journal of Roentgenology, 151(1), 171–179.

Bassett, D. S., Yang, M., Wymbs, N. F., & Grafton, S. T. (2015). Learning-induced autonomy of sensorimotor systems. Nature Neuroscience, 18(5), 744.

Behrens, T. E. J., Johansen-Berg, H., Woolrich, M. W., Smith, S. M., Wheeler-Kingshott, C. A. M., Boulby, P. A., … Matthews, P. M. (2003). Non-invasive mapping of connections between human thalamus and cortex using diffusion imaging. Nature Neuroscience, 6(7), 750. http://doi.org/10.1038/nn1075

Bell, P. T., & Shine, J. M. (2016). Subcortical contributions to large-scale network communication. Neuroscience & Biobehavioral Reviews, 71, 313–322. http://doi.org/http://dx.doi.org/10.1016/j.neubiorev.2016.08.036

Birney, & Bowman. (2009). An experimental-differential investigation of cognitive complexity. Psychology Science Quarterly, 51(4), 449–469. Retrieved from http://p16277.typo3server.info/fileadmin/download/PschologyScience/4-2009/psq_4_2009_449-469.pdf

Birney, D. P., Halford, G. S., & Andrews, G. (2006). Measuring the Influence of Complexity on Relational Reasoning: The Development of the Latin Square Task. Educational and Psychological Measurement, 66(1), 146–171. http://doi.org/10.1177/0013164405278570

Bollmann, S., Puckett, A. M., Cunnington, R., & Barth, M. (2018). Serial correlations in single-subject fMRI with sub-second TR. NeuroImage, 166, 152–166.

Braun, U., Plichta, M. M., Esslinger, C., Sauer, C., Haddad, L., Grimm, O., … Meyer-Lindenberg, A. (2012). Test-retest reliability of resting-state connectivity network characteristics using fMRI and graph theoretical measures. NeuroImage, 59(2), 1404–1412. http://doi.org/10.1016/j.neuroimage.2011.08.044

Brett, M., Anton, J.-L., Valabregue, R., & Poline, J.-B. (2002). Region of interest analysis using an SPM toolbox.

Chao-Gan, Y., & Yu-Feng, Z. (2010). DPARSF: A MATLAB toolbox for “pipeline” data analysis of resting-state fMRI. Frontiers in Systems Neuroscience, 4(13), 1–7. http://doi.org/10.3389/fnsys.2010.00013

Cocchi, L., Halford, G. S., Zalesky, A., Harding, I. H., Ramm, B. J., Cutmore, T., … Mattingley, J. B. (2014). Complexity in relational processing predicts changes in functional brain network dynamics. Cerebral Cortex, 24(9), 2283–2296. http://doi.org/10.1093/cercor/bht075

Cocchi, L., Zalesky, A., Fornito, A., & Mattingley, J. B. (2013). Dynamic cooperation and competition between brain systems during cognitive control. Trends in Cognitive Sciences, 17(10), 493–501.

Cole, M. W., Bassett, D. S., Power, J. D., Braver, T. S., & Petersen, S. E. (2014). Intrinsic and task-evoked network architectures of the human brain. Neuron, 83(1), 238–251. http://doi.org/10.1016/j.neuron.2014.05.014

Cole, M. W., & Schneider, W. (2007). The cognitive control network: Integrated cortical regions with dissociable functions. NeuroImage, 37(1), 343–60. http://doi.org/10.1016/j.neuroimage.2007.03.071

Crittenden, B. M., Mitchell, D. J., & Duncan, J. (2016). Task encoding across the multiple demand cortex is consistent with a frontoparietal and cingulo-opercular dual networks distinction. Journal of Neuroscience, 36(23), 6147–6155. http://doi.org/10.1523/JNEUROSCI.4590-15.2016

Damle, N. R., Ikuta, T., John, M., Peters, B. D., DeRosse, P., Malhotra, A. K., & Szeszko, P. R. (2017). Relationship among interthalamic adhesion size, thalamic anatomy and neuropsychological functions in healthy volunteers. Brain Structure and Function, 222(5), 2183–2192. http://doi.org/10.1007/s00429-016-1334-6

Desikan, R. S., Ségonne, F., Fischl, B., Quinn, B. T., Dickerson, B. C., Blacker, D., … Hyman, B. T. (2006). An automated labeling system for subdividing the human cerebral cortex on MRI scans into gyral based regions of interest. Neuroimage, 31(3), 968–980.

Dhollander, T., Raffelt, D., & Connelly, A. (2016). Unsupervised 3-tissue response function estimation from single-shell or multi-shell diffusion MR data without a co-registered T1 image. In ISMRM Workshop on Breaking the Barriers of Diffusion MRI (Vol. 5).

Dosenbach, N., Fair, D. A., Miezin, F. M., Cohen, A. L., Wenger, K. K., Dosenbach, R. A. T., … Petersen, S. E. (2007). Distinct brain networks for adaptive and stable task control in humans. Proceedings of the National Academy of Sciences, 104(26), 11073–11078. http://doi.org/10.1073/pnas.0704320104

Dosenbach, N., Visscher, K. M., Palmer, E. D., Miezin, F. M., Wenger, K. K., Kang, H. C., … Petersen, S. E. (2006). A core system for the implementation of task sets. Neuron, 50(5), 799–812. http://doi.org/10.1016/j.neuron.2006.04.031

Duncan, J. (2010). The multiple-demand (MD) system of the primate brain: mental programs for intelligent behaviour. Trends in Cognitive Sciences, 14(4), 172–9. http://doi.org/10.1016/j.tics.2010.01.004

Duquette, M., Rainville, P., Alary, F., Lassonde, M., & Lepore, F. (2008). Ipsilateral cortical representation of tactile and painful information in acallosal and callosotomized subjects. Neuropsychologia, 46(8), 2274–2279. http://doi.org/10.1016/j.neuropsychologia.2008.02.017

Eklund, A., Nichols, T. E., & Knutsson, H. (2016). Cluster failure: why fMRI inferences for spatial extent have inflated false-positive rates. Proceedings of the National Academy of Sciences, 113(28), 7900–7905. http://doi.org/10.1073/pnas.1602413113

Frazier, J. A., Chiu, S., Breeze, J. L., Makris, N., Lange, N., Kennedy, D. N., … Dieterich, M. E. (2005). Structural brain magnetic resonance imaging of limbic and thalamic volumes in pediatric bipolar disorder. American Journal of Psychiatry, 162(7), 1256–1265.

Friston, K. J., Holmes, A. P., Worsley, K. J., Poline, J., Frith, C. D., & Frackowiak, R. S. J. (1994). Statistical parametric maps in functional imaging: a general linear approach. Human Brain Mapping, 2(4), 189–210.

Gazzaniga, M. S., Bogen, J. E., & Sperry, R. W. (1962). Some functional effects of sectioning the cerebral commissures in man. Proceedings of the National Academy of Sciences of the United States of America, 48, 1765–1769. http://doi.org/10.1073/pnas.48.10.1765

Glass, H. C., Shaw, G. M., Ma, C., & Sherr, E. H. (2008). Agenesis of the corpus callosum in California 1983-2003: A population-based study. American Journal of Medical Genetics, Part A, 146(19), 2495–2500. http://doi.org/10.1002/ajmg.a.32418

Goldstein, J. M., Seidman, L. J., Makris, N., Ahern, T., O’Brien, L. M., Caviness, V. S., … Tsuang, M. T. (2007). Hypothalamic abnormalities in schizophrenia: sex effects and genetic vulnerability. Biological Psychiatry, 61(8), 935–945.

Halford, Wilson, & Phillips, S. (1998). Processing capacity defined by relational complexity: implications for comparative, developmental, and cognitive psychology. The Behavioral and Brain Sciences, 21(6), 803–31; discussion 831-64. Retrieved from http://www.ncbi.nlm.nih.gov/pubmed/10191879

Hearne, L. J., Cocchi, L., Zalesky, A., & Mattingley, J. B. (2015). Interactions between default mode and control networks as a function of increasing cognitive reasoning complexity. Human Brain Mapping, 36(7), 2719–2731. http://doi.org/10.1002/hbm.22802

Hearne, L. J., Cocchi, L., Zalesky, A., & Mattingley, J. B. (2017). Reconfiguration of brain network architectures between resting-state and complexity-dependent cognitive reasoning. The Journal of Neuroscience, 37(35), 12–13. http://doi.org/10.1111/ijlh.12426

Hetts, S. W., Sherr, E. H., Chao, S., Gobuty, S., & Barkovich, a J. (2006). Anomalies of the corpus callosum: an MR analysis of the phenotypic spectrum of associated malformations. AJR. American Journal of Roentgenology, 187(5), 1343–8. http://doi.org/10.2214/AJR.05.0146

Hinkley, L. B. N., Marco, E. J., Brown, E. G., Bukshpun, P., Gold, J., Hill, S., … Nagarajan, S. S. (2016). The Contribution of the Corpus Callosum to Language Lateralization. Journal of Neuroscience, 36(16), 4522–4533. http://doi.org/10.1523/JNEUROSCI.3850-14.2016

Honey, C. J., Sporns, O., Cammoun, L., Gigandet, X., Thiran, J. P., Meuli, R., & Hagmann, P. (2009). Predicting human resting-state functional connectivity from structural connectivity. Proceedings of the National Academy of Sciences of the United States of America, 106(6), 2035–40. http://doi.org/10.1073/pnas.0811168106

Hwang, K., Bertolero, M. A., Liu, W. B., & D’Esposito, M. (2017). The human thalamus is an integrative hub for functional brain networks. Journal of Neuroscience, 37(23), 5594–5607.

Jeurissen, B., Tournier, J.-D., Dhollander, T., Connelly, A., & Sijbers, J. (2014). Multi-tissue constrained spherical deconvolution for improved analysis of multi-shell diffusion MRI data. NeuroImage, 103, 411–426.

Lassonde, M., Sauerwein, H., Chicoine, A. J., & Geoffroy, G. (1991). Absence of disconnexion syndrome in callosal agenesis and early callosotomy: Brain reorganization or lack of structural specificity during ontogeny? Neuropsychologia, 29(6), 481–495. http://doi.org/10.1016/0028-3932(91)90006-T

Makris, N., Goldstein, J. M., Kennedy, D., Hodge, S. M., Caviness, V. S., Faraone, S. V, … Seidman, L. J. (2006). Decreased volume of left and total anterior insular lobule in schizophrenia. Schizophrenia Research, 83(2), 155–171.

Marco, E. J., Harrell, K. M., Brown, W. S., Hill, S. S., Jeremy, R. J., Kramer, J. H., … Paul, L. K. (2012). Processing speed delays contribute to executive function deficits in individuals with agenesis of the corpus callosum. Journal of the International Neuropsychological Society□: JINS, 18(3), 521–9. http://doi.org/10.1017/S1355617712000045

Mišić, B., Betzel, R. F., De Reus, M. A., van den Heuvel, M. P., Berman, M. G., McIntosh, A. R., & Sporns, O. (2016). Network-level structure-function relationships in human neocortex. Cerebral Cortex, 26(7), 3285–3296. http://doi.org/10.1093/cercor/bhw089

Owen, J. P., Li, Y.-O., Yang, F. G., Shetty, C., Bukshpun, P., Vora, S., … Mukherjee, P. (2013). Resting-state networks and the functional connectome of the human brain in agenesis of the corpus callosum. Brain Connectivity, 3(6), 547–62. http://doi.org/10.1089/brain.2013.0175

Parrish, M. L., Roessmann, U., & Levinsohn, M. W. (1979). Agenesis of the corpus callosum: a study of the frequency of associated malformations. Annals of Neurology: Official Journal of the American Neurological Association and the Child Neurology Society, 6(4), 349–354.

Paul, L. K., Brown, W. S., Adolphs, R., Tyszka, J. M., Richards, L. J., Mukherjee, P., & Sherr, E. H. (2007). Agenesis of the corpus callosum: genetic, developmental and functional aspects of connectivity. Nature Reviews. Neuroscience, 8(4), 287–99. http://doi.org/10.1038/nrn2107

Paul, L. K., Erickson, R. L., Hartman, J. A., & Brown, W. S. (2016). Learning and memory in individuals with agenesis of the corpus callosum. Neuropsychologia, 86, 183–192. http://doi.org/10.1016/j.neuropsychologia.2016.04.013

Paul, L. K., Schieffer, B., & Brown, W. S. (2004). Social processing deficits in agenesis of the corpus callosum: narratives from the Thematic Appreciation Test. Archives of Clinical NeuropsychologyLJ: The Official Journal of the National Academy of Neuropsychologists, 19(2), 215–25. http://doi.org/10.1016/S0887-6177(03)00024-6

Putnam, M. C., Wig, G. S., Grafton, S. T., Kelley, W. M., & Gazzaniga, M. S. (2008). Structural organization of the corpus callosum predicts the extent and impact of cortical activity in the nondominant hemisphere. The Journal of Neuroscience□: The Official Journal of the Society for Neuroscience, 28(11), 2912–8. http://doi.org/10.1523/JNEUROSCI.2295-07.2008

Quigley, M., Cordes, D., Turski, P., Moritz, C., Haughton, V., Seth, R., & Meyerand, M. E. (2003). Role of the corpus callosum in functional connectivity. American Journal of Neuroradiology, 24(2), 208–212. http://doi.org/10.1016/0014-4886(87)90083-5

Roland, J. L., Snyder, A. Z., Hacker, C. D., Mitra, A., Shimony, J. S., Limbrick, D. D., … Leuthardt, E. C. (2017). On the role of the corpus callosum in interhemispheric functional connectivity in humans. Proceedings of the National Academy of Sciences, 114(50), 13278–13283. http://doi.org/10.1073/pnas.1707050114

Romaniello, R., Marelli, S., Giorda, R., Bedeschi, M. F., Bonaglia, M. C., Arrigoni, F., … Borgatti, R. (2017). Clinical characterization, genetics, and long-term follow-up of a large cohort of patients with agenesis of the corpus callosum. Journal of Child Neurology, 32(1), 60–71. http://doi.org/10.1177/0883073816664668

Schmitt, L. I., Wimmer, R. D., Nakajima, M., Happ, M., Mofakham, S., & Halassa, M. M. (2017). Thalamic amplification of cortical connectivity sustains attentional control. Nature, 1–24. http://doi.org/10.1038/nature22073

Schulte, T., & Müller-Oehring, E. M. (2010). Contribution of callosal connections to the interhemispheric integration of visuomotor and cognitive processes. Neuropsychology Review, 20(2), 174–190.

Seeley, W. W., Menon, V., Schatzberg, A. F., Keller, J., Glover, G. H., Kenna, H., … Greicius, M. D. (2007). Dissociable intrinsic connectivity networks for salience processing and executive control. The Journal of Neuroscience: The Official Journal of the Society for Neuroscience, 27(9), 2349–56. http://doi.org/10.1523/JNEUROSCI.5587-06.2007

Seymour, S. E., Reuter-lorenz, P. A., & Gazzaniga, M. S. (1994). The disconnection syndrome: Basic findings reaffirmed. Brain, 117(1), 105–115. http://doi.org/10.1093/brain/117.1.105

Shen, K., Mišić, B., Cipollini, B. N., Bezgin, G., Buschkuehl, M., Hutchison, R. M., … Berman, M. G. (2015). Stable long-range interhemispheric coordination is supported by direct anatomical projections. Proceedings of the National Academy of Sciences, 112(20), 6473–6478. http://doi.org/10.1073/pnas.1503436112

Siffredi, V., Anderson, V., Leventer, R. J., & Spencer-Smith, M. M. (2013). Neuropsychological profile of agenesis of the corpus callosum: a systematic review. Developmental Neuropsychology, 38(February 2015), 36–57. http://doi.org/10.1080/87565641.2012.721421

Smith, R. E., Tournier, J.-D., Calamante, F., & Connelly, A. (2012). Anatomically-constrained tractography: improved diffusion MRI streamlines tractography through effective use of anatomical information. Neuroimage, 62(3), 1924–1938.

Smith, R. E., Tournier, J. D., Calamante, F., & Connelly, A. (2013). SIFT: Spherical-deconvolution informed filtering of tractograms. NeuroImage, 67, 298–312. http://doi.org/10.1016/j.neuroimage.2012.11.049

Sun, F. T., Miller, L. M., & D’Esposito, M. (2004). Measuring interregional functional connectivity using coherence and partial coherence analyses of fMRI data. NeuroImage, 21(2), 647–658. http://doi.org/http://dx.doi.org/10.1016/j.neuroimage.2003.09.056

Tomasch, J. (1954). Size, distribution, and number of fibres in the human corpus callosum. The Anatomical Record, 119(1), 119–135.

Tournier, J. D., Calamante, F., & Connelly, A. (2010). Improved probabilistic streamlines tractography by 2nd order integration over fibre orientation distributions. In Proc 18th Annual Meeting of the Intl Soc Mag Reson Med (ISMRM) (p. 1670).

Tournier, J. D., Calamante, F., & Connelly, A. (2012). MRtrix: Diffusion tractography in crossing fiber regions. International Journal of Imaging Systems and Technology, 22(1), 53–66. http://doi.org/10.1002/ima.22005

Tovar-Moll, F., Moll, J., de Oliveira-Souza, R., Bramati, I., Andreiuolo, P. a, & Lent, R. (2007). Neuroplasticity in human callosal dysgenesis: a diffusion tensor imaging study. Cerebral Cortex (New York, N.Y.: 1991), 17(3), 531–41. http://doi.org/10.1093/cercor/bhj178

Tovar-Moll, F., Monteiro, M., Andrade, J., Bramati, I. E., Vianna-Barbosa, R., Marins, T., … Lent, R. (2014). Structural and functional brain rewiring clarifies preserved interhemispheric transfer in humans born without the corpus callosum. Proceedings of the National Academy of Sciences of the United States of America, 111(21), 7843–7848. http://doi.org/10.1073/pnas.1400806111

Tyszka, J. M., Kennedy, D. P., Adolphs, R., & Paul, L. K. (2011). Intact bilateral resting-state networks in the absence of the corpus callosum. The Journal of Neuroscience: The Official Journal of the Society for Neuroscience, 31(42), 15154–62. http://doi.org/10.1523/JNEUROSCI.1453-11.2011

Urbanski, M., Bréchemier, M.-L., Garcin, B., Bendetowicz, D., Thiebaut de Schotten, M., Foulon, C., … Volle, E. (2016). Reasoning by analogy requires the left frontal pole: lesion-deficit mapping and clinical implications. Brain, 139(6), 1783–1799. http://doi.org/10.1093/brain/aww072

van der Knaap, L. J., & van der Ham, I. J. M. (2011). How does the corpus callosum mediate interhemispheric transfer? A review. Behavioural Brain Research, 223(1), 211–221.

Wahl, M., Strominger, Z., Jeremy, R. J., Barkovich, a J., Wakahiro, M., Sherr, E. H., & Mukherjee, P. (2009). Variability of homotopic and heterotopic callosal connectivity in partial agenesis of the corpus callosum: a 3T diffusion tensor imaging and Q-ball tractography study. AJNR. American Journal of Neuroradiology, 30(2), 282–9. http://doi.org/10.3174/ajnr.A1361

Woolgar, A., Parr, A., Cusack, R., Thompson, R., Nimmo-Smith, I., Torralva, T., … Duncan, J. (2010). Fluid intelligence loss linked to restricted regions of damage within frontal and parietal cortex. Proceedings of the National Academy of Sciences of the United States of America, 107(33), 14899–902. http://doi.org/10.1073/pnas.1007928107

Yeo, B. T., Krienen, F. M., Sepulcre, J., Sabuncu, M. R., Lashkari, D., Hollinshead, M., … Buckner, R. L. (2011). The organization of the human cerebral cortex estimated by intrinsic functional connectivity. Journal of Neurophysiology, 106(3), 1125–1165. http://doi.org/10.1152/jn.00338.2011.

Zalesky, A., Fornito, A., & Bullmore, E. T. (2010). Network-based statistic: identifying differences in brain networks. NeuroImage, 53(4), 1197–207. http://doi.org/10.1016/j.neuroimage.2010.06.041

